# The geometry of appetitive-aversive value representations in medial prefrontal networks

**DOI:** 10.1101/2023.03.03.530871

**Authors:** Nanci Winke, Cyril Herry, Daniel Jercog

**Author notes:** These authors contributed equally to this work (Cyril Herry, Daniel Jercog).

## Abstract

The value of rewards and punishments – namely, how good or bad they are perceived – guides approach or avoidance behaviors. Valence refers to the negative or positive “sign” of the state elicited by an event, whereas salience refers to the amount of attention an event attracts, disregarding its valence. While identifying these signals conveys critical information for understanding circuits involved in emotional processing, they are often confounded due to their underlying correlation. Moreover, whereas the study of the neural basis of value coding has been intensively investigated in the appetitive domain, the neural substrates for how aversive values are established for different threat intensities and guide defensive behavior have yet to be discovered. The dorsomedial prefrontal cortex (dmPFC) is a key region in the control of defensive actions, although how different aversive values are encoded at the neuronal level within this region and drive defensive behaviors remains unknown.

Here, we developed an instrumental approach/avoidance task in mice that, by matching motivational salience levels elicited by cues predicting rewards or punishments, allows univocally disentangling the presence of either salience, valence, or value coding from brain signals. We performed freely moving large neuronal population calcium imaging in the dmPFC of mice performing our task, conducting appetitive/aversive outcome devaluation/revaluation behavioral tests. We found that, while a similar fraction of single neurons decoded valence and value information, and only a minor fraction decoded salience, value coding was observed at the neuronal population level. Moreover, different value representations of the same valence lay within similar subspaces of the neural state space while values of opposed valence were encoded in orthogonal subspaces, unveiling how the brain stores associative appetitive and aversive information in medial prefrontal networks.

## Introduction

The value of predictive cues is a fundamental feature that guides behavior. A common approach to the study of the substrates of value signals is by analyzing preferences during choice behavior: choosing one option over another reveals the preference elicited by a particular item or stimulus (Rangel et al., 2008; O’Doherty 2014). While such an approach may have clear advantages in revealing the actual value on which choices are based, active choice might engage decision processes that entangle their information with value signals. In contrast, an alternative approach from behavioral neuroscience defines value in terms of the motivating properties of a stimulus for an instrumental action, where the degree to which an animal is prepared to perform some effortful action to obtain/avoid a stimulus relates to the degree to which the animal finds that stimulus rewarding/aversive (O’Doherty 2014; Rolls, 2007).

While value quantifies how good or bad something is, salience signals the amount of attention that captures. Salience and value are strongly correlated in the reward domain — a more rewarding outcome has a higher value (it is more desirable), as well as higher salience (as it is more important) — whereas salience and value are negatively correlated in the domain of punishments. In contrast, valence can be considered as signaling a positive or negative affect, according to whether the stimulus is associated with dislike or pleasure (Tye, 2018).

The dorsomedial prefrontal cortex (dmPFC) in mice, can be defined as comprising the cingulate cortex (Cg1) and prelimbic area (PL) (Laubach, 2018; but see Le Merre et al., 2021). The dmPFC is critical in adaptive behavior and executive control (Dalley et al., 2004), and is a central structure in the regulation of defensive actions (Jercog et al., 2021), however, how different aversive values are represented within this area and influence defensive behavior remains unknown. Here we investigated i) how different threat intensities are encoded in dmPFC networks, ii) what information aspect, regarding salience, valence or value information, is encoded when representing appetitive-aversive outcome predicting cues within the dmPFC. Using a novel behavioral paradigm that allows the univocally disentangling of such information in freely-moving mice while performing large population calcium-imaging recordings, we found that the dmPFC populations encode value information for appetitive-aversive predicted outcomes. Moreover, aversive and appetitive values representations by dmPFC populations were encoded in orthogonal manner.

## Results

To first assess how information of cues associated to different threat intensities is represented in medial prefrontal networks, mice were initially submitted to a differential avoidance conditioning task. In this task, mice were placed in a shuttlebox consisting in 2 symmetric compartments separated by a small hurdle. After an inter-trial interval, and independently of their current location, they are challenged with one of three possible auditory stimuli (CS): CSs and CSw, paired at 7 sec after onset with a strong- and weak-footshock, respectively; and CSn, which was neutral (**Figure 1a**, **Supplementary Figure 1**). Moreover, moving away from the current compartment (shuttling) during a CS presentation before 7 sec defined an avoidance response, which stopped the ongoing auditory CS and prevented any contingent shock (**Figure 1a**). After learning, mice shuttled with a rate proportional to the magnitude of the shock size predicted by the CSs, CSw, respectively **(Figure 1b**, **Supplementary Figure 1**). Additionally, CSs and CSw induced different vigor in avoidance responses (**Figure 1c**).

**Figure 1.**
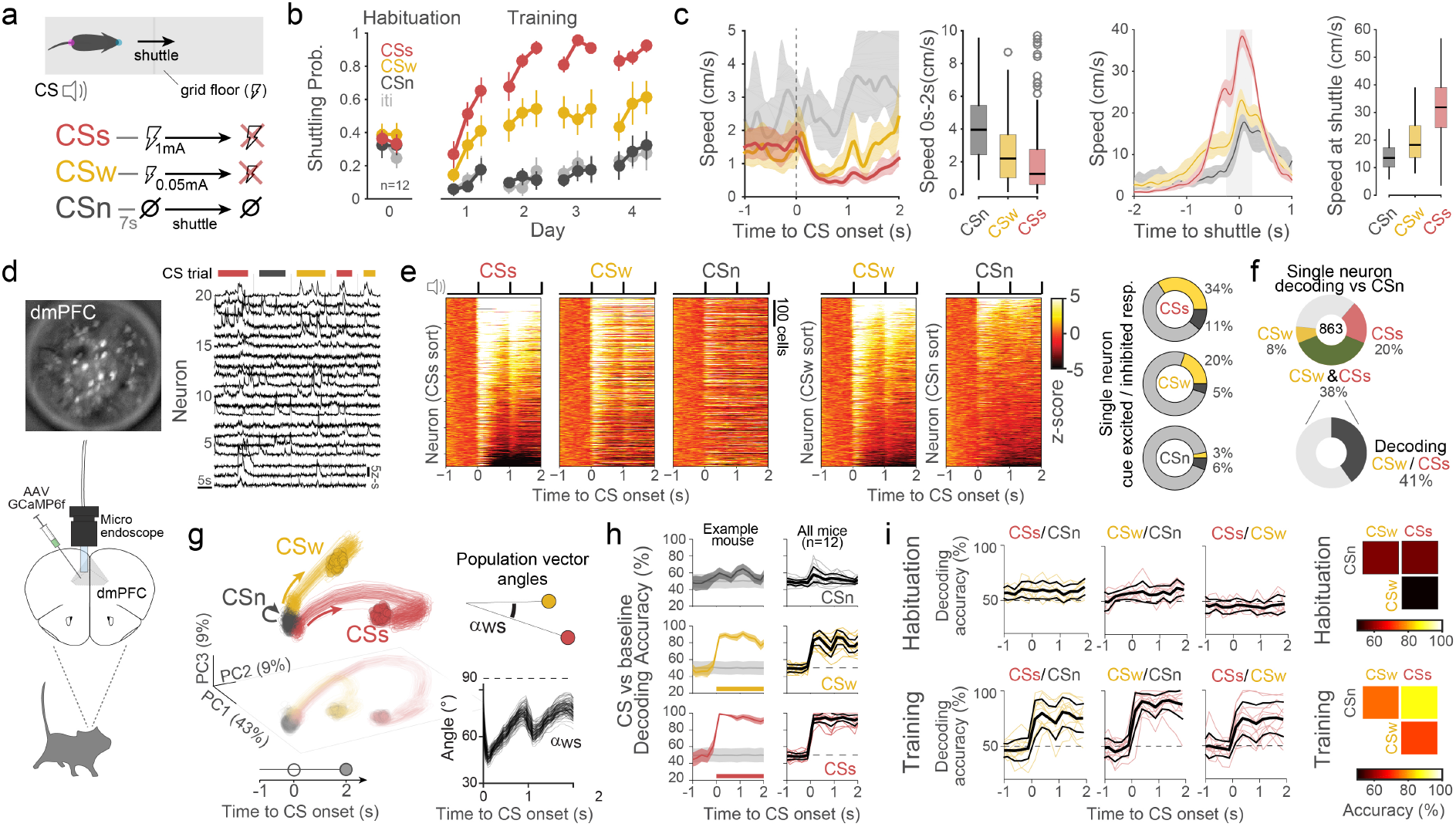
Different threat intensity coding in dorsomedial prefrontal networks. **a.** Schematics of the behavioral task. After an inter-trial interval and independently of the location of the animal, an auditory CS starts. Shuttling away from the current compartment before 7 sec stops the ongoing CS and prevents any contingent footshock (avUS). CSs and CSw, are associated with two different avUS intensities (1mA and 0.05mA, respectively), whereas CSn is not associated with any outcome. CS were presented in random order. **b.** Shuttle probability for different trial type. Significantly different shuttling probabilities between CS-types from the third 10-trial block on Day 1 (P < 0.05; Two-way RM ANOVA; n = 12 mice). **c.** CS onset-triggered (left panel) and compartment-crossing (shuttle) triggered (right panel) median speed for CSn, CSw, and CSs shuttling trials (shuttle latency >3 s). **d.** dmPFC calcium-imaging population recordings in freely behaving mice. GCaMP6m was genetically expressed in CamKII neurons, and a GRIN lens was implanted targeting the dmPFC. Calcium fluorescence traces of 20 example neurons for 5 trials (red, CSs; grey, Csn; yellow, CSw). **e.** Z-scored activity for CSs, CSw, and CSn ordered by CSs responses magnitude (left; Day 4: 12 mice, n = 863 neurons). Z-scored activity for CSw and CSn is ordered by their self-response magnitude (center). Percentage of total neurons significantly excited (yellow), inhibited (dark gray), or not modulated (light gray) for each CS (right) **f.** Fraction of single neurons significantly decoding CSw or CSs from CSn (top; **Methods**). Fraction of subpopulation significantly decoding CSw from CSs (bottom). **g.** Neural trajectories for CS-onset triggered averaged responses for CSn, CSw, and CSs, projected onto the first 3 principal components (average responses over 50% of trials, 100 repetitions) (left). Arrows schematize trajectory direction. Instantaneous population vector angles between CSw and CSs response (right) **h.** dmPFC population activity-based decoding accuracies at single mouse level for each CS from baseline. Representative mouse decoding accuracies (left). Shuffled trial label in gray. Accuracy data display mean +/- SD. Thick lines indicate significant decoding accuracies (P < 0.05, Permutation test). Individual mice mean decoding accuracies (right). Median +/- MAD in black. **i.** Left, Single-mouse dmPFC population activity-based decoding accuracies for CS identity during habituation (top) and training day 4 (bottom). Right, Median matrix for mean decoding accuracies across mice (0 to 2 s after CS onset) for habituation (top) and training day 4 (bottom).

The medial prefrontal cortex is a key brain region in the control of instrumental defensive actions (Diehl et al., 2018; Capuzzo Floresco, 2020; Jercog et al., 2021). To compare how different associated threat intensities are represented in medial prefrontal networks, we expressed the calcium-indicator GCaMP6f under the CaMKII promoter in the dorsomedial prefrontal cortex (dmPFC) of 12 wild-type mice via a GRIN lens targeting this area (**Figure 1d, Supplementary Figure 2, Methods**). Training induced more prevalent neuronal responses to CSs and CSw compared to CSn, and while many of these neurons displayed increased activity in response to threat-predictive cues, other neurons exhibited/inhibitory responses (**Figure 1e**, **Supplementary Figure 2**). Moreover, while 38% of neurons decoded threat-predicting cues from CSn, 41% from this subpopulation also decoded threat identity (**Figure 1f**). Despite that CSs and CSw induced a significant global activation of dmPFC populations (**Figure 1e**), the encoding of both threat-predicting stimuli did not reflect a proportionality on the same ensemble’s response, as it is shown in the behavior (**Figure 1b**). Instead, CSw and CSs engaged different neuronal ensembles, expressed by their divergent neural population low dimensional trajectories and vector angles between elicited population responses (**Figure 1g, Supplementary Figure 2**). To quantify differences in neuronal population representations for different threat intensities in dmPFC networks, we used a population decoding approach using machine learning-based linear classifiers for individual animals (Jercog et al., 2021). By decoding CS from baseline activity, we first observed that learning induced sustained representations for both CSw and CSs (**Figure 1g, Supplementary Figure 3**). In addition, CSn was weakly represented by dmPFC populations in agreement with our previous observations (Jercog et al., 2021). Moreover, CSs and CSw representations were induced by training and consistently differ across recorded animals (**Figure 1h, Supplementary Figure 3**). Overall, cues associated with different threat intensities elicited different population representations and dynamics in dmPFC networks.

We observed that different CS-associated threat intensities elicited distinct representations in dmPFC networks. Previous human studies on emotional processing have accounted that dissociating salience, valence, and value information from brain signals requires tasks including the appetitive and aversive domains, to assess how similarly strong/weak positive and negative outcomes are represented (Litt et al., 2011; Kahnt et al., 2014; Zhang et al., 2017) (**Figure 2a**). In the same line, we hypothesize that if animals can perform the same action to approach rewards as avoiding punishments, and this is achieved at the same level of performance for a reward-predicting cue as for one of the threat-predicting cues, i.e. eliciting the same motivational salience, it allows to unequivocally disentangle if neural representations encode salience, valence or value (**Figure 2b**). Thus, in a second experiment, after training on the differential avoidance task, mice (n=10) were water restricted and submitted to an instrumental appetitive protocol to learn that shuttling from the current compartment during the presentation of another auditory cue (CSr) lead to water rewards (**Figure 2c**, **Supplementary Figure 4, Methods**). Animals learned to discriminate the four different randomly presented auditory cues while exhibiting not significantly different levels of shuttling to avoid during CSs or to approach during CSr (**Figure 2d**). In addition, although CSs and CSr also induced similar strength on associated conditioned shuttle responses (**Supplementary Figure 5**), CSr induced ballistic responses towards reward ports in contrast to CSs shuttling, showing that mice discriminated CSs from CSr from a behavioral perspective (**Figure 2e**).

**Figure 2.**
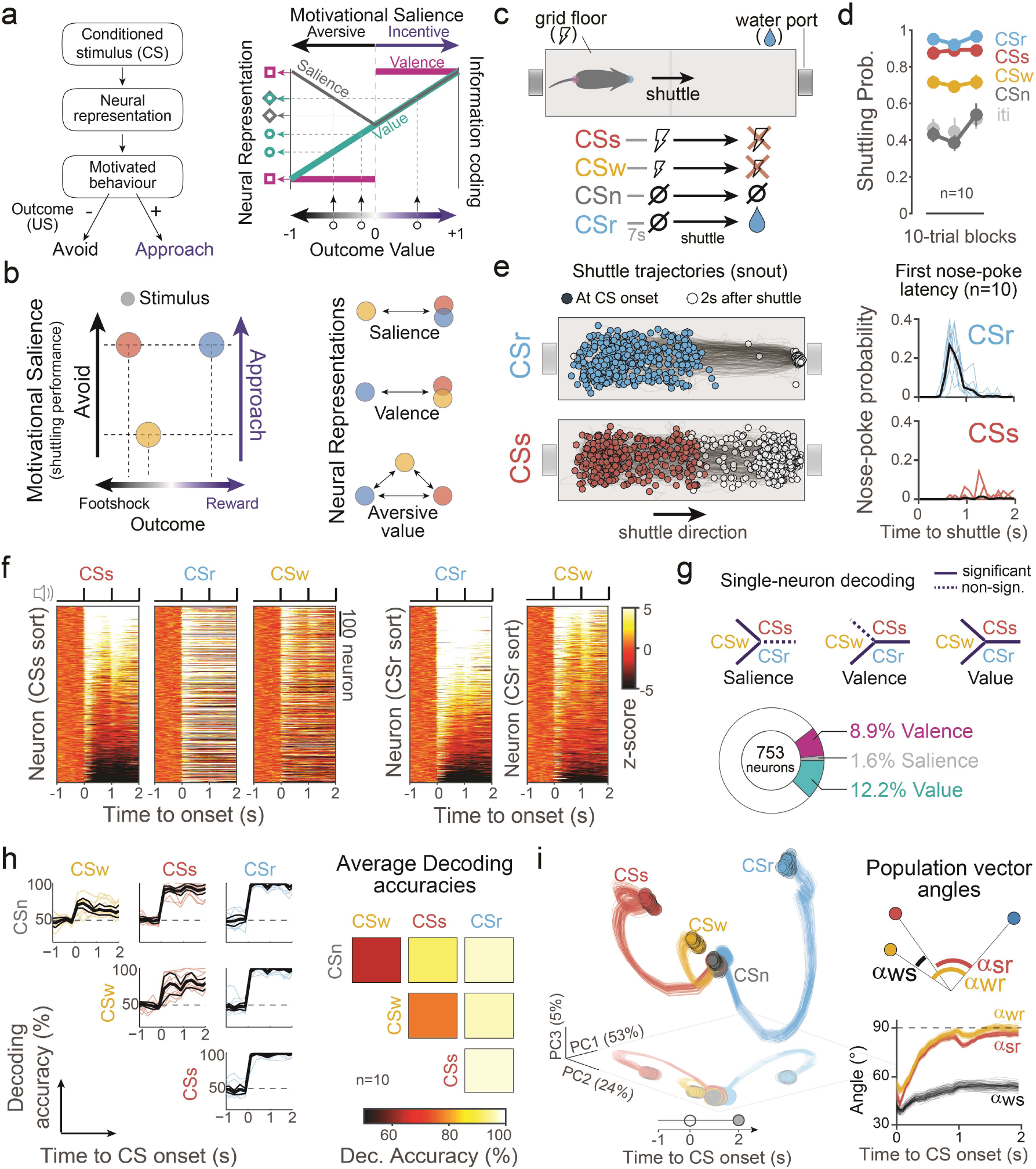
dmPFC neuronal population represents value coding. **a.** A given event associated with a certain outcome, such as a CS, leads to neuronal representations that promote motivated behaviors. If the associated outcome is aversive or appetitive, motivated behavior elicited will be avoidance or approach, respectively (left). Neural representations within a brain region can convey different emotional processing aspects related to the internal value of an associated outcome: i) salience representations (gray), reflecting the magnitude of outcome values disregarding their positive or negative sign, ii) valence representations (purple), reflecting the negative or positive sign of outcome values, disregarding their magnitude, iii) value representation (green), reflecting magnitude and sign of outcome values (right). Notice that three chosen outcome values (black circles) can lead to 3 different representations if the region encodes value information (green circles), whereas only 2 different representations would be elicited if encoding salience (gray diamonds) or valence (purple squares). **b.** Two stimuli eliciting the same magnitude of incentive (water reward) and aversive (footshock) salience, reflected by equal shuttling performance, and another stimulus eliciting a different level of motivational salience, allows disentangling if associated neural representations encode salience, valence or value. **c.** Instrumental approach/avoidance task, adding to the previous task the CSr associated with a water reward. Water-restricted mice gain access to the reward by shuttling from the current compartment to the CSr before 7 s. **d.** Shuttle probabilities for different CS-type during calcium imaging recording session. Different shuttle probabilities for CSw and CSs/r (P<0.05) and non-significantly probabilities between CSs and CSr (P = 0.47) (Two-way RM ANOVA; n = 10 mice). **e.** Shuttle trajectories for CSs and CSr trial types. Colored circles indicate the animal’s location at CS onset, and white circles the location 2 s after the shuttle (left). Nosepoke probability for CSs and CSr trial types (right). **f.** Z-scored activity for CSs, CSr, and CSw ordered by CSs responses magnitude (left). Z-scored activity for CSr and CS was ordered by their self-response magnitude (right) (n = 10 mice, n = 753 neurons). **g.** Single-neuron decoding accuracies (see **Methods**). A salience neuron significantly decodes CSw from CSs and CSr but not CSs from CSr. A valence neuron significantly decodes CSr from CSs and CSw but not CSs from CSw. A value neuron decodes all pairs of CS. **h.** Individual-mouse-based dmPFC populations decoding accuracies at CS onset. Each line represents the dmPFC population activity-based decoding accuracies for an individual mouse. Median +/- MAD decoding accuracies are overlaid in black (left). Median matrix for mean decoding accuracies from 0 to 2 s (right). **i.** Neuronal population trajectories for the four different trial types projected onto the first 3 PC for CS onset-aligned conditions (left). Population vector angles between CSw and CSs (gray, ws), CSw and Csr (yellow, wr), and CSs and CSr (red, sr).

As for CSs and CSw, CSr elicited responses by dmPFC neurons, globally exhibiting a larger fraction of excited neurons compared to the aversive CS (**Figure 2f, Supplementary Figure 5**). While a similar fraction of single neurons decoded valence and value information (8.9% and 12.2%, respectively), a smaller fraction of neurons decoded salience (1.6%) (**Figure 2g**). Moreover, we observed significant decoding between CS representations at the neuronal population level, consistent with value coding in the dmPFC (**Figure 2h, Supplementary Figure 6**). Notably, low dimensional dynamics and angles between population vectors showed that appetitive and aversive predictive cues’ information were encoded by dmPFC neuronal population activity in an orthogonal fashion, where population vector angles between CSs and CSw were decreased (**Figure 2i, Supplementary Figure 6**).

To further investigate how changes in appetitive and aversive values associated with the same stimuli were represented by dmPFC populations, we next exposed animals to an incentive devaluation protocol where mice performed the same task but after non-restricted (*ad libitum*) access to water in their home cages (**Figure 3a**). After the devaluation, mice maintained the same shuttling levels for aversive CS, whereas CSr shuttling dropped to similar levels as for CSw (**Figure 3a**). In this scenario, CSr evoked population activity became more similar to the one elicited by all the other stimuli, as decoding accuracies from CSr to other stimuli were reduced, particularly to CSw (**Figure 3b**). Notably, the encoding of the devalued CSr still preserved the orthogonal directionality to the trajectories induced by the aversive CSs and CSw (**Figure 3c**).

**Figure 3.**
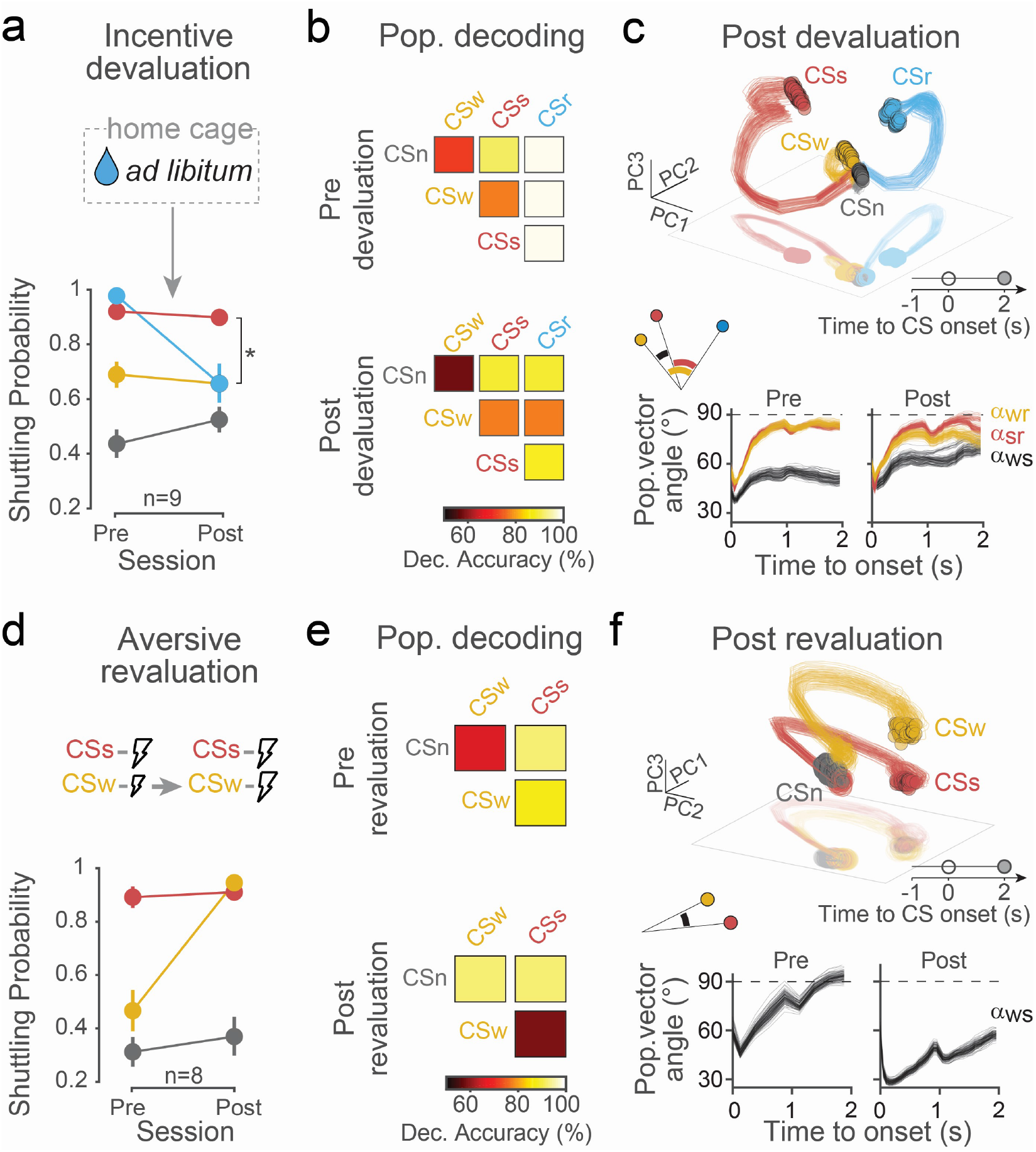
Effect of behavioral manipulations on the dmPFC neuronal population. **a.** Incentive devaluation, after a regular session of instrumental approach/avoidance training, mice were given *ad libitum* water access in their home cage and tested the following day. The shuttle probability for CSr after incentive devaluation significantly decreased (CSr vs CSs: Pre, P = 0.3671; Post, P < 0.05) to the same levels as CSw (CSr vs CSw: Pre, P < 0.05; Post, p > 0.99) (Two-way RM ANOVA; n = 9 mice). **b.** Median matrix for the mean decoding accuracies of single-mouse for pre (top) and post-devaluation (bottom). After the devaluation, decoding accuracy for CSr from other types of CS decreases. **c.** Neuronal population trajectories for the four different trial types projected onto the first 3 principal components, for CS onset-aligned conditions (top). Population vector angles (bottom) between CSw and CSs (gray, ws), CSw and Csr (yellow, wr) and CSs and CSr (red, sr). **d.** Aversive revaluation, animals were first submitted to a regular differential avoidance conditioning followed by a session in which the US intensity for both stimuli was the one typically associated with the CSs. After revaluation training, shuttle probability for the CSw matched one of the CSs (CSw vs CSs: Pre, P < 0.05; Post, P = 0.6) (Two-way RM ANOVA; n = 8 mice). **e.** Median matrix for mean decoding accuracies of single-mouse for pre (top) and post-revaluation (bottom). After the revaluation, decoding accuracies between CSw and CSs decrease, whereas between CSw and CSn increases. **f.** Neuronal population trajectories for the three different trial types projected onto the first 3 principal components (top). Population vector angle between CSw and CSs (gray, ws).

Finally, to assess how different threat-predicting cues with same outcomes are represented in the dmPFC, we next performed an aversive revaluation protocol where CSw became associated with the same footshock outcome associated with CSs (**Figure 3d**). Training progressively induced an increase in shuttling for CSw towards the same levels as CSs (**Figure 3d**). In this scenario, the ability to decode CSw and CSs from dmPFC population activity was largely diminished (**Figure 3e**). Consistent with this observation, low dimensional trajectories for CSw and CSs became similar as population vector angles were largely reduced (**Figure 3f**) suggesting that the information encoded at CS onset mainly reflects information about the associated outcome. Altogether, these findings are consistent with the dmPFC populations representing the value of appetitive and aversive associated cues, where opposed valence representations are encoded in an orthogonal manner.

## Discussion

In this study, by performing calcium-imaging recordings in the dmPFC of mice and characterizing the neural population representations of appetitive and aversive stimuli associated with different outcome magnitudes, we found that dmPFC populations encode value information of appetitive and aversive associations. Using a different set of linear classifiers and population analyses, we showed that dmPFC neuronal populations encode the value of appetitive-aversive outcome-associated cues. Moreover, we showed that different values of same-valence were overall encoded within subspaces that were orthogonal to those encoding values of opposed valence. Notably, the core of these findings observed with calcium-imaging approaches were reproduced by using electrophysiological single-unit recording techniques (**Supplementary Figure 2, 3** and **6**). In addition, these electrophysiological experiments confirmed that sustained representations observed in calcium-imaging data were not due to slow calcium indicator kinetics and were, instead, consistent with our previous observations of sustained threat representations in the dmPFC (Jercog et al., 2021). Altogether, our results reveal how the information of different values of appetitive and aversive predictive cues are represented in medial prefrontal networks.

Identifying true value signals requires including different magnitudes for both positive and negative outcomes in the experimental designs, which has previously been accounted for by human studies (Kahnt et al., 2014; Litt et al., 2011; Zhang et al., 2017). Following this principle, we developed a novel behavioral task in mice where animals learn that different cues predict reward and punishment outcomes associated with the performance of an equivalent instrumental shuttling response. A similar approach was used to assess valence coding by using a single appetitive and aversive associated outcome (Kyriazi et al., 2018). Our experimental framework relies on the assumption that approach/avoidance probabilities are a proxy to individuals’ values of associated outcomes. Thus, by eliciting the same level of shuttling for the reward and one of the two punishment-predicting cues, this matching in motivational salience allows us to univocally determine the presence of salience, valence and value information from brain signals. Thus, this minimal set of stimuli configuration can account for studying true value signals using mice.

In the context of associative learning, previous rodent studies have described valence coding in regions including the basolateral nucleus of the amygdala (Beyeler et al., 2016; Zhang Li, 2018, Kyriazi et al., 2018) and medial prefrontal networks (Kyriazi et al,. 2020). While some of these studies use different or the presence-or-absence of conditioned responses as a readout of motivational salience, using the same elicited conditioned responses during the presentation of appetitive and aversive predicting cues (Kyriazi et al., 2018, 2020) allows us to rule out the possibility that impending divergent motor responses influence differences in encoding of appetitive and aversive predicting cues. In addition, while in some of these previous studies the conditioned responses overlaps with the conditioned stimuli presentation likely entangling the stimulus and response encoding of information, our analyses were restricted to CS onset periods before the initiation of shuttling responses (Jercog et al., 2021). Decoding conditioned stimuli associated with positive and negative outcomes while matching the motivational salience of conditioned responses allows to rule out possible salience confounding, allowing to determine if stimuli are perceived as negative or positive (Beyeler et al., 2016; Kyriazi et al., 2020). Further, our framework showed that addressing the dissociation of value from valence information requires eliciting different levels of motivational salience, hence the presence of at least an additional conditioned stimuli in the appetitive/aversive domain.

We found that representations of associated stimuli of opposed valence were encoded in an orthogonal manner by the activity of dmPFC populations (**Figure 2i**), consistent with the idea that opposed nonoverlapping ensembles of neurons in dmPFC networks largely drive opposed valence associative representations. Despite changes in the similarity of representations when devaluating reward outcomes, such orthogonal geometry in the encoding of information of opposed valence associations was still preserved (**Figure 3c**). Notably, the amygdala has accounted for such information encoding structure to signal opposed valence (Kyriazi et al., 2018; Grundemann et al., 2019). In addition, while learning is reported to induce an increase in similarity by a reduction of angles of population patterns between conditioned stimuli and aversive reinforcers in the amygdala (Grewe et al., 2017), we observed that the presence or absence of the reward-associated stimulus strikingly influenced such similarity measurements between prefrontal aversive associative representations. Indeed, although CSw and CSs outcome contingencies were not changed, population vector angles between CSw and CSs representations (**Figure 1g**) were diminished during the avoid-approach task (**Figure 2i**), while reward devaluation increased them (**Figure 3c**). In addition, removing the reward component before the aversive revaluation procedure reinstated maximal population vector angles between CSw and CSs representations (**Figure 3f**). These results are in line with previous observations in decision-making contexts, where aversive values were encoded in a relative coding rather than in an absolute coding manner (Rangel and Clithero, 2012).

In addition, changing the aversive outcome associated with CSw to the one associated with CSs, neural population patterns elicited by both stimuli became highly similar (**Figure 3e**). While this shows that dmPFC associative representations mainly do not represent sensory properties of stimuli, it suggests that representations are linked to the value of expected outcomes. Indeed, we have previously shown that preventing avoidance responses towards a threat-predicting stimulus while changing its contingencies in a similar paradigm largely preserves representations of threat-predicting stimuli, suggesting that threat representations at CS onset are not reflecting information about the impending defensive actions (Jercog et al., 2021). Further experiments will be required on the appetitive domain to address whether this is a general principle that disregards the valence nature of associations, or if different outcome modalities eliciting either the same aversive or incentive salience can elicit similar representations within dmPFC networks.

The ability to decode the different appetitive and aversive associated stimuli (**Figure 2-3**) supports that the dmPFC encodes value information of associations. Recent fMRI human studies on value coding, controlling for possible salience confounds, agree on the ACC as encoding for salience information (Kahnt et al., 2014; Litt et al., 2011; Zhang et al., 2017). The dmPFC in mice includes the frontal cingulate cortex area, which is suggested to be homologous to the human ACC (van Heukelum et al., 2020). This is in contrast to our observation of dmPFC encoding values, supported also by the fact that only a minor fraction of neurons encoded salience compared to value or valence information (**Figure 2g**). Since fMRI signals for the average activation within voxel areas, one possible explanation for the discrepancy with our results could be the inability of fMRI to identify different neuronal ensembles eliciting a similar global activation within a voxel (Zhang et al., 2017). Indeed, our results suggest this scenario, as averaging the activity across the monitored ensembles increases the similarity in representations to the extent that CSs and CSr become indistinguishable after reward devaluation, which would be accounted as salience coding (**Supplementary Figure 7**). Thus, our findings suggest revising the underlying coding of the salience network defined by fMRI studies. Future experiments will be necessary to address the precise coding scheme from other brain regions.

Finally, psychiatric conditions such as anxiety and obsessive-compulsive disorders are associated with overestimating the costs and probabilities of threats (Sookman and Pinard 2002; Peschard and Philippot, 2017; Grupe and Nitschke, 2013). The dmPFC is a central structure where cognitive and motivational control converges with emotional generation and regulation (Kober et al., 2008). Here we showed that dmPFC populations orthogonally represent aversive and appetitive values of associations. The origin of value signals has been a matter of intense research. The orbitofrontal cortex (OFC) is primarily attributed to signaling reward values to drive decision processes (Rangel et al., 2008; Knudsen and Wallis, 2022), broadcasting this information to different areas including dorsolateral prefrontal (Wallis Miller, 2003) and anterior cingulate (Kennerley and Wallis, 2009; Enel et al., 2020) cortices. Whether there is a common circuitry leading to the origin of appetitive and aversive value signals is still debated (Levy and Schiller, 2021). Notably, simultaneous recordings in the amygdala and OFC of non-human primates reveal that, while positive valence-coding neurons tend to shift representations of stimulus-outcome contingencies faster in the OFC, negative valence-coding neurons shift representations faster in the amygdala, suggesting that the underlying circuitry signaling values might be different for appetitive and aversive conditions (Morrison et al., 2011a,b). Consistent with this idea of different circuits for appetitive and aversive processing, stimuli of opposite valence engage different ensembles in the amygdala, likely activating different downstream projections (Beyeler et al., 2016; Zhang et al., 2021). The medial prefrontal cortex is an integration area that is critical in adaptive behavior and executive control (Dalley et al., 2004) (O’Doherty 2011). We previously reported that the dmPFC has a critical role in instrumental defensive behavior through a dynamic process of linking threat representations with those associated with initiating defensive actions (Jercog et al., 2021). In this scenario, what is the precise role of the aversive values coding in the dmPFC and how it affects the dynamic process of selection of defensive actions, is a matter of future research.

## Methods

### Subject detail

Adult male C57BL6/J mice (Janvier) of 8-20 weeks old were individually housed under a 12 h light-dark cycle and provided with food and water ad libitum. All procedures were performed per standard ethical guidelines (European Communities Directive 86/60-EEC) and were approved by the Animal Health and Care committee of Institut National de la Santé et de la Recherche Médicale and French Ministry of Agriculture and Forestry (agreement #A3312001). Prior to training in the instrumental approach/avoidance task, mice were placed on water restriction while ensuring that their body weight remained at ≥ 85% of initial values. Water restriction consisted of a maximum of 15 consecutive days of restriction. Mice were handled and habituated to be connected for three days before the experiment started. All data correspond to implanted and connected animals.

### Behavioral apparatus

The differential avoidance conditioning task and the instrumental approach/avoidance task were performed in a shuttlebox comprised of a plexiglass box (40 x 10 x 30 cm) with a floor grid connected to a shocker, where a small plastic hurdle (1 cm height) divided the arena into two equal compartments while infrared beams detection automatically monitored the mice shuttling between compartments (Imetronic). On both extremities of the shuttlebox, a water port with a white LED mounted internally, a photogate, and a steel drinking tube was placed 2 cm above the grid floor (SanWorks). A programmable controller (Arduino, Spyder) controlled LED activation and water delivery through a fast solenoid valve connected to a water container. The shuttlebox was enclosed inside an acoustic foam isolated box where two speakers mounted on top of each compartment delivered the auditory conditioned stimulus (CS) consisting in either 3 kHz, 7 kHz, 13 kHz, or white-noise 50 ms pips at 1 Hz (maximum 13 pips, i.e., CS maximum duration of 12 s), 2 ms rise and fall, 80 dB sound pressure level. Scrambled foot shocks of 50 ms (~7 Hz) at an intensity of either 1 mA or 0.05 mA and a maximum duration of 5 s applied through the grid floor (Imetronic) served as the aversive unconditioned stimulus (avUSstrong or avUSweak, respectively). Water delivery of 20 μL by the drinking tube served as the appetitive unconditioned stimulus (apUS, Extended Data Figure 1). LEDs mounted on top of each compartment provided house light. During the Confined task, one opaque white wall was placed in the middle of the maze, and another additional wall blocked the water port preventing shuttling between compartments and access to the water delivery. A video camera recorded from above at 30 fps for video-tracking purposes using Open Broadcaster Software Studio (OBS, version 26.0.2). For the subset of mice used for electrophysiological recordings (Extended Data Figure x), another video camera recorded from above at 30/40 fps for video-tracking purposes using Cineplex Software (Cineplex, Plexon).

### Behavioral paradigm

#### Differential avoidance conditioning task

There were three different auditory stimuli (CS) in this task: the CSs, consisted of 3 kHz pips and paired with the avUSstrong, the CSw, consisted of 7 kHz pips and paired with the avUSweak; and the CSn, comprised of white-noise pips that were neutral and therefore not paired with the US. Mice were first habituated to the context and tones (Day 0). During the habituation, the three CS were presented 20 times in an intermingled fashion without any US. Training days (Day > 0) consisted of randomly intermingling 30 CSs, 30 CSw and 30 CSn, where CSs and CSw trials were paired with the avUSstrong and avUSweak, respectively if mice did not shuttle before 7 s after CS onset. The CS started independently from the animal location in the shuttlebox after a random inter-trial interval of 25-40 s and shuttling between compartments stopped any ongoing CS or US. The first CSs and CSw trial of the first training session was always paired with the US disregarding the behavior of the animal. All sessions started with 2 min acclimation periods before the first CS presentation. During training, mice were connected with miniaturized fluorescence microscope dummies except for the habituation (Day 0) and the last training day (Day 4), where calcium imaging recordings were performed using a miniaturized fluorescence microscope (nVoke, Inscopix). For more details, see the section below, Calcium imaging recordings.

#### Instrumental approach/avoidance task

Mice (n=10) were first submitted to the differential avoidance conditioning task and then placed under water restriction protocol. In addition to the CSs, CSw and CSn, a new auditory stimulus was introduced. The CSr consisted of 13 kHz pips paired with an apUS. Training consisted of randomly intermingling 50 CSs, 50 CSw, 50 CSr and 50 CS. As before, the CS started independently from the animal location in the shuttlebox after an inter-trial interval of 15-20 s, and shuttling between compartments stopped any ongoing CS. For the CSs and CSw, shuttling prevented the aversive US, whereas for the CSr shuttling gave access to the appetitive US. A nose poke on the port of the shuttled compartment before 12 s of the onset of CSr triggers the delivery of the water reward. Mice were trained with miniaturized fluorescence microscope dummies until the shuttle probability for the CSs and CSr significantly matched, upon which calcium imaging recordings were performed using a miniaturized fluorescence microscope (nVoke, Inscopix). For more details, see the section below, Calcium imaging recordings.

#### Reward outcome devaluation training

Mice were then submitted to a training session of instrumental approach/avoidance task as described in the previous section. Following instrumental training, reward outcome devaluation was achieved by allowing animals to *ad libitum* water access in their home cage 24h before training. Calcium imaging recordings were collected during training sessions before and after the devaluation of the reward outcome.

#### Aversive outcome revaluation training

Following instrumental training, aversive outcome revaluation was accomplished by changing the outcome associated with the CSw. The outcome that before was the avUSweak of 0.05 mA was now replaced with the avUSweak of 1 mA. As such, both CSw and CSs have the same outcome. Mice were submitted to at least three aversive outcome revaluation training sessions. The training stopped once the shuttle probability between CSw and CSs matched. Calcium imaging recording sessions were performed the day before, the first and the last day of the aversive outcome revaluation.

### Surgical procedure

Mice (8-10 weeks old) were anesthetized with isoflurane (induction 3%, maintenance 1. 5%) in O2. For analgesia, a subcutaneous injection of 0.05 mL of metacam (5 mg/kg body weight) was administered 30 min before anesthesia. Additionally, 0.1 mL of local lidocaine anesthesia (Lidor, 20 mg/mL diluted with sodium chloride at 0.5%) was applied under the scalp before the incision. Mice were placed in a stereotaxic frame (Kopf Instruments), body temperature was maintained at 37°C with a temperature controller system (FHC), and eyes were hydrated with Lacrigel (Europhta Laboratories). All implants were secured using 3 stainless steel screws placed before the craniotomy.

### Virus injection and GRIN lens implantation

Following craniotomy, 200 μL of GCaMP6f (AAV5.CamKII.GCaMP6f.WPRE.SV40, titer 2.3×1013, University of PennsylvaniaVector Core) was unilaterally injected into the dmPFC using a micromanipulator (Scientifica) and pulled glass pipettes (tip diameter ~25 mm) at the following coordinates from bregma: + 2.1 mm AP; - 0.5 mm ML; and 1.20 mm DV from dura with an angle of 6 degrees. After injection, a track above the imaging site was opened with a sterile needle (26G, 0.45 mm outer diameter, Terumo) to assist in inserting the GRIN lens (0.5 x 4.0 mm, Inscopix). The needle was inserted at the exact coordinates of the injection, except for the DV, at 1.0 mm. The GRIN lens was subsequently lowered by the micromanipulator at a speed of 1μm/s through the track until 1.26 mm from the brain surface. Super Bond cement was used to secure the GRN lens and to make the base structure necessary for the baseplate once optimal GCaMP expression was achieved. Mice were allowed to recover for at least 4 weeks after surgery before checking for GCaMP expression.

### Calcium imaging recordings

Four to six weeks after surgery, mice were anesthetized with isoflurane to fix the miniature microscope baseplate on top of the cranium implant using dental cement. The miniature microscope was detached, and the baseplate was covered with a cover. One week before starting the behavioral training, mice were habituated to the mounting procedure and the weight of the miniscope for four consecutive days. For the first three days, animals were restrained, the baseplate cover removed, a miniature microscope dummy was attached to the baseplate, and mice were left with it for at least one hour in their home cage. On the last day, mice were mounted with the miniature microscope to set the miniature microscope light-emitting diode (LED) power and electronic focus. Imaging data were acquired using the nVoke software at a frame rate of 20 Hz (exposure time, 50 ms) with a LED power of 0.5 to 0.8 mW/mm2, an analogue gain of 8 and a spatial downsampling by a factor of 2. The recording sessions were not recorded in a continuous way. Instead, we recorded in segments around the CS presentation. Each segment consisted of recording for 3 sec before the CS onset and 2 sec after the end of the trial. For the CSs and CSw, the end of the trial was considered the shuttle before 7 sec or after the escape of the aversive outcome. For the CSn, the end of the trial was considered the shuttle before 7 sec or until the end of the CSn. For the CSr, the end of the trial was after water delivery or at the end of the CSr. Therefore, each segment lasts between 5 to 17 sec. Inputs triggered by Arduino automatically controlled the start and end of each recorded segment. Events such as CS onset, shuttle and nose pokes were recorded in the Inscopic software acquisition by TTL inputs.

### Processing of calcium imaging data

We preprocessed the data using the Inscopix Data Processing Software (IDPS). Given the high number of trials per session, up to 200 trials, all trials had to be merged into a time series, exported into a movie and then uploaded again for the remaining preprocessing. We used spatial downsampling of factor by 2, cropped the field-of-vire to only the lens area, and used the spatial filter, motion correction and F/F with the default settings. We then proceeded to the identification of the neurons using a PCA-ICA analysis. We manually validated each neuron by inspecting its morphology, activity trace and calcium transients. Individual deltaF/F calcium traces were detrended and z-scored for further calculations.

### Electrode implantation and electrophysiological recordings

Following craniotomy, mice were bilaterally implanted in the dmPFC with the electrode array targeting the following coordinates relative to bregma: + 2.0-2.1 mm AP; ± 0.55-0.7 mm ML; and 1.20-1.30 mm DV from dura with an angle of 14 degrees. Each electrode bundle consisted of 16 individually insulated nichrome wires (13 mm diameter, impedance 60–100 KU; Kanthal) fixed to an electrode guide. Each electrode bundle was attached to one 18-pin connector (Omnetics). Connectors were referenced and grounded via silver wires (127 μm diameter, A-M Systems) placed above the cerebellum. All implants were secured using Super-Bond cement (Sun Medical). After surgery, mice were allowed to recover for at least 10 days. Electrodes were connected to a headstage (Plexon) containing sixteen unity-gain operational amplifiers. Each headstage was connected to a 16-channel PBX preamplifier where the signal was replicated and bandpass-filtered at 300 Hz and 8 kHz and at 0.5 Hz and 200 Hz for local field potential recordings. Spiking activity was digitized at 40 kHz and isolated by time-amplitude window discrimination and template matching using an Omniplex system (Plexon). Single-unit spike sorting was performed using Off-Line Spike Sorter (OFSS, Plexon), where Pairwise-P (multivariate ANOVA) statistics were used to assess unit isolation quality (P < 0.05). At the end of the experiment, electrolytic lesions were administered before transcardial perfusion to verify electrode tip location using standard histological techniques.

### Histological analyses

Mice were administered a lethal dose of Exagon with lidocaine and underwent transcardial perfusions via the left ventricle with 4% w/v paraformaldehyde (PFA) in 0.1 M PB. In electrode-implanted animals, electrolytic lesions of 10 μA during 5 s(Stimulus Isolator, WPI) to identify electrode bundle tip location were performed prior to PFA perfusions. Following dissection, brains were post-fixed for 24 h at 4°C in 4% PFA. For GRIN lense implanted animals, the head was left post-fixed for 24 h at 4°C in 4% PFA and only then dissected. Brain sections of 100 μm-thick were cut on a vibratome, mounted on gelatin-coated microscope slides, and dried. To verify viral injections and GRIN location in dmPFC, serial 100 μm-thick slices containing the regions of interest were mounted in VectaShield (Vector Laboratories) and imaged using an epifluorescence system (Leica DM 5000) fitted with a 10 x dry objective.

The location of the electrode bundle tip or GRIN lenses was visually inspected. Only recordings targeting the dmPFC were included in the analyses. The location and the extent of the injections/infections were visually controlled. Only infections targeting the dmPFC were included in the analyses.

### Data Analysis

Single neuron instantaneous activity was computed in 100ms bins (50ms sliding window) and average across trials for heatmaps were z-scored relative to the last second before CS onset. We use principal component analysis for dimensionality reduction and data visualization purposes, by considering 80% of trials to get estimates of single neuron mean activity, repeating the procedure 100 times. This same data was used to assess the population vector (PV) angles, which instead were computed in the full neural space. Angles between PV for different trial types A and B, following:

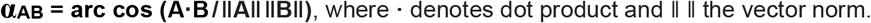

#### Decoding analyses

We trained support-vector machine linear classifiers for single-neuron and single-mouse based population decoding. Decoding was performed independently in 250ms binned activity (see Jercog et al., 2021 for full description). The decoding accuracy of a classifier was defined as the proportion of correctly classified trials following a 5-fold cross validation procedure. The estimate of the mean accuracy was obtained by averaging the results from repeating the procedure 500 times. Shuffle accuracies were estimated by randomly shuffling the label the classes over 1000 repetitions. Significant decoding was defined when mean accuracy exceeded those from the null distribution provided by the shuffle accuracies, following a permutation test (Jercog et al., 2021). Decoding analyses from electrophysiological recordings was performed by constructing pseudo-population vectors containing the joint activity from different mice (n = 4) as described in (Jercog et al., 2021), where individual spike counts were binned in 0.5 s non-overlapping windows.

Snout tracking was obtained using DeepLabCut (Mathis et al., 2018) (network initially trained with 100 uniformly distributed frames in 5 videos, one extra iteration to correct outlier detections).

We used custom routines written in MATLAB (The MathWorks) to perform all data analyses.

### Statistics

Bonferroni-correction post hoc tests were used for ANOVA multiple comparisons if a significant main effect or interaction was observed. Box plots indicate median (center), interquartile range (box) and most extreme data points (whiskers) that were not considered outliers (1.5× the interquartile range away from the box, in circles). Statistical analyses were performed with Matlab (MathWorks) and PRISM (GraphPad Software). All behavioral experiments were controlled by computer systems, and data were collected in an automated and unbiased way. No statistical methods were used to predetermine sample sizes, but sample sizes were based on previous studies in our laboratory. Mice that did not satisfy the active avoidance learning criteria (CSs avoidance probability greater than 0.3 at day 3) or decreasing below 85% weight during water restriction (appetitive phase) were discarded from the study. No other mice or data points were excluded.

## Acknowledgements

We thank members of the Herry Laboratory for helpful discussions. S. Laumond, J. Tessaire and the technical staff of the housing and experimental animal facility of the Neurocentre Magendie. This work was supported by grants from the French National Research Agency (ANR719 DOPAFEAR, ANR-PRELONGIN, ANR-10-EQPX-08 OPTOPATH), the Conseil Regional d’Aquitaine, and the Fondation pour la Recherche Médicale (FRM-Equipes FRM 2017).

## Supplementary Figures

**Supplementary Figure 1.**
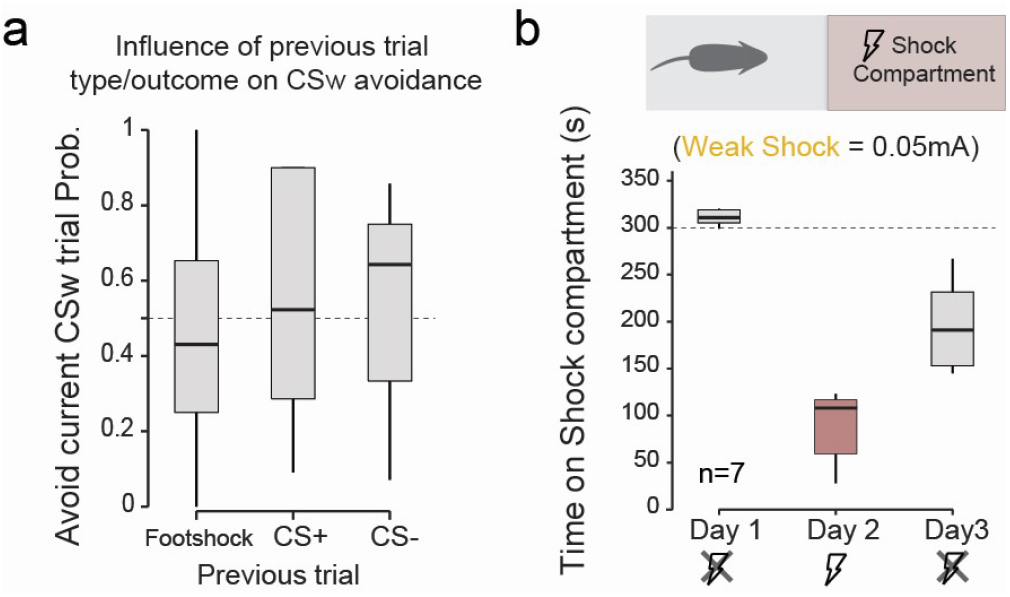
**a**. Influence on previous trial type/outcome on current CSw avoidance. **b**. Place-preference induced by weak shock (0.05 mA, associated to CSw in the differential avoidance task). Animals receive electric footshock as they enter and stay in the shock compartment, and footshock stopped upon moving away from it.

**Supplementary Figure 2.**
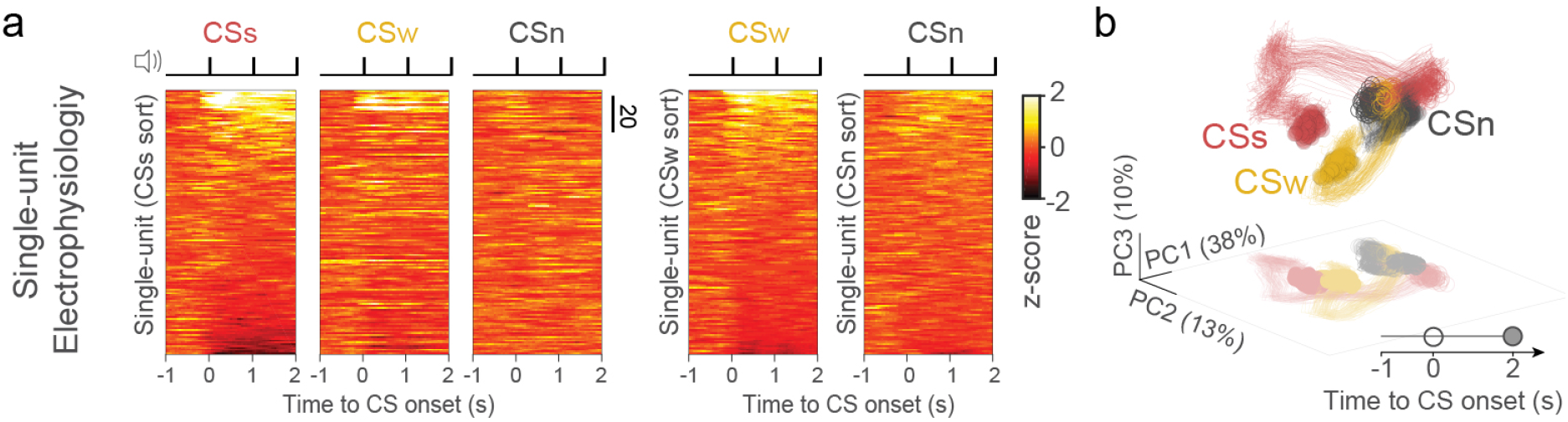
Electrophysiological single-unit recordings in the dmPFC during the differential avoidance conditioning task (Day 4). **a.** Z-scored activity for CSs, CSw, and CSn ordered by CSs responses magnitude (left; Day 4: 4 mice, n = 145 units). Z-scored activity for CSw and CSn is ordered by their self-response magnitude (right). Notice the different level of activation induced by the CS presentation, explained by the fact that for electrophysiological recordings, single-unit identified can be activated during inter-trial intervals (missing in our calcium-imaging to avoid possible bleaching). **b**. Neural trajectories for CS-onset triggered averaged responses for CSn, CSw, and CSs, projected onto the first 3 principal components (average responses over 80% of trials, 100 repetitions).

**Supplementary Figure 3.**
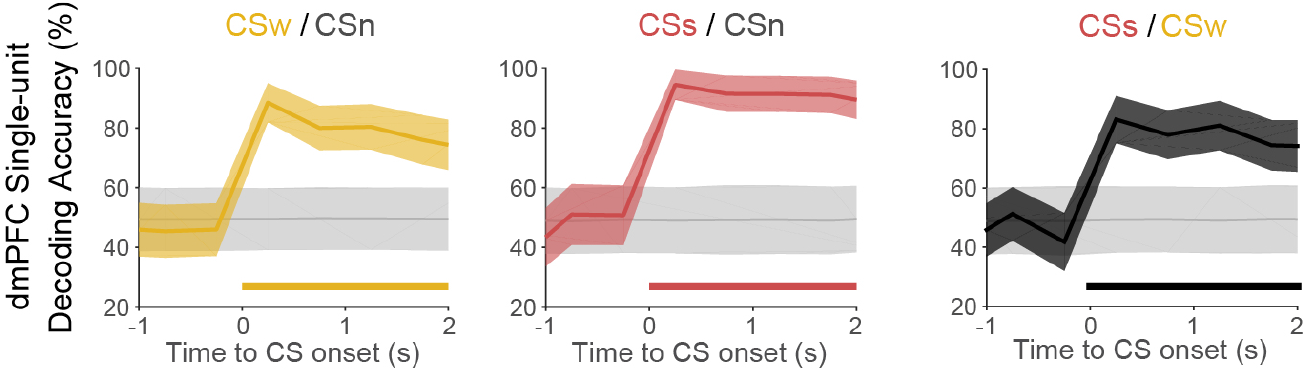
Decoding accuracy at CS onset from electrophysiological single-unit pseudo-populations (Day 4; 4 mice, 145 neurons). Pseudo-population consists of the different single-units recorded across mice as in (Jercog et al., 2021). CSw from CSn (left), CSs from CSn (center), CSs from CSw (right). Accuracy data display mean +/- SD. Thick lines indicate significant decoding accuracies (P < 0.05, Permutation test).

**Supplementary Figure 4.**
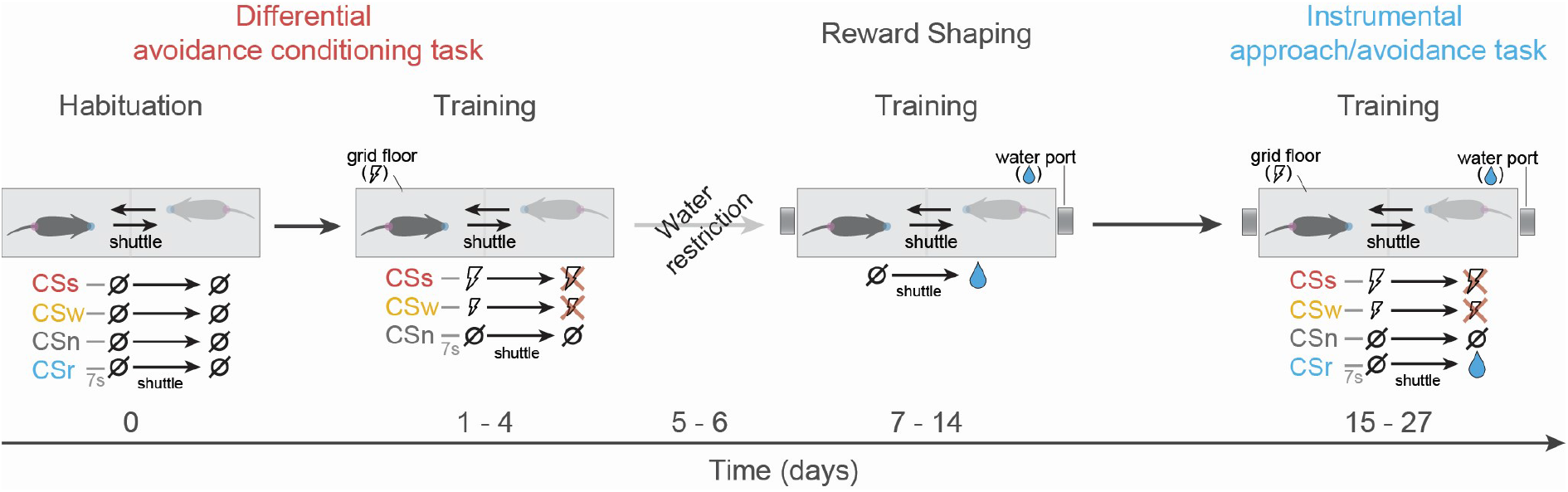
Protocol for the instrumental approach/avoidance task. Mice are initially submitted to a differential avoidance conditioning task. At day 0, mice undergo a habituation session in which four different auditory stimuli are presented 20 times in an intermingled fashion (CSstrong – 3 kHz; CSweak – 7 kHz; CSneutral – white noise; CSreward – 13 kHz). The differential avoidance conditioning task training consists of two auditory stimuli associated with two different footshock intensities and a neutral auditory stimulus. After an inter-trial interval and independently of the animal’s location, an auditory CS starts. Shuttling away from the current compartment before 7 sec stops the ongoing CS and prevents any contingent footshock (avUS). CSs and CSw, are associated with two different avUS intensities (1mA and 0.05mA, respectively), whereas CSn is not associated with any outcome. Each CS is presented 30 times and presented in random order. Following differential avoidance training, mice are water-restricted for two days, receiving 1 mL daily in their home cage. At reward shaping, mice progressively learn that shuttling from the current compartment gains them access to a water reward after nose poking. The duration of the shaping depends on each mouse. Once a mouse reaches 80% of performance, they are moved to the following task. The shaping has a minimum duration of 4 days and a maximum of 7 days, after which, if they do not perform well, the behavior terminates, and mice are given water in their home cage. In the last stage of the protocol, a new association is presented in which CSr is associated with a water reward. Water-restricted mice gain access to the reward by shuttling from the current compartment to the CSr before 7 s. On the first two days of the training, the CS are presented in blocks: a reward block consisting of CSr and CSn and an aversive block consisting of CSs, CSw and CSn. Each block is presented twice and contains 25 Cs of each type. The session starts with an appetite block. Subsequently, the training consists of 50 presentations of each CS, presented intermingled.

**Supplementary Figure 5.**
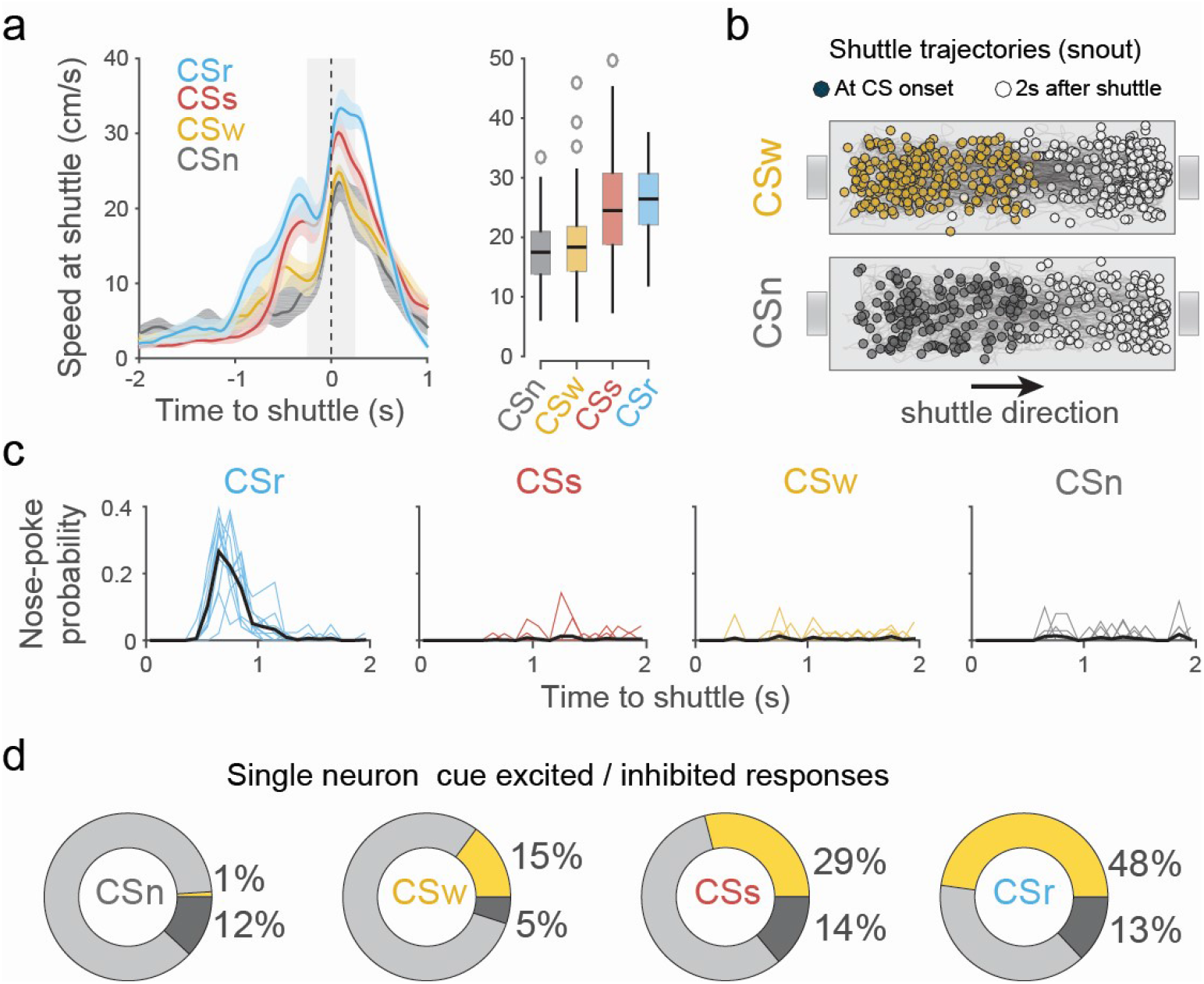
Approach/avoidance task. **a.** Shuttle triggered median speed for CSn, CSw, CSs and CSr (shuttle latency >3 s). **b**. Shuttle trajectories for CSw and CSn trial types. Colored circles indicate the animal’s location at CS onset, and white circles the location 2 s after the shuttle. **c**. Nose-poke probability after shuttle during a CS for different trial types. **d**. Percentage of total cells significantly excited (yellow), inhibited (dark gray), or not modulated (light gray) for each CS type.

**Supplementary Figure 6.**
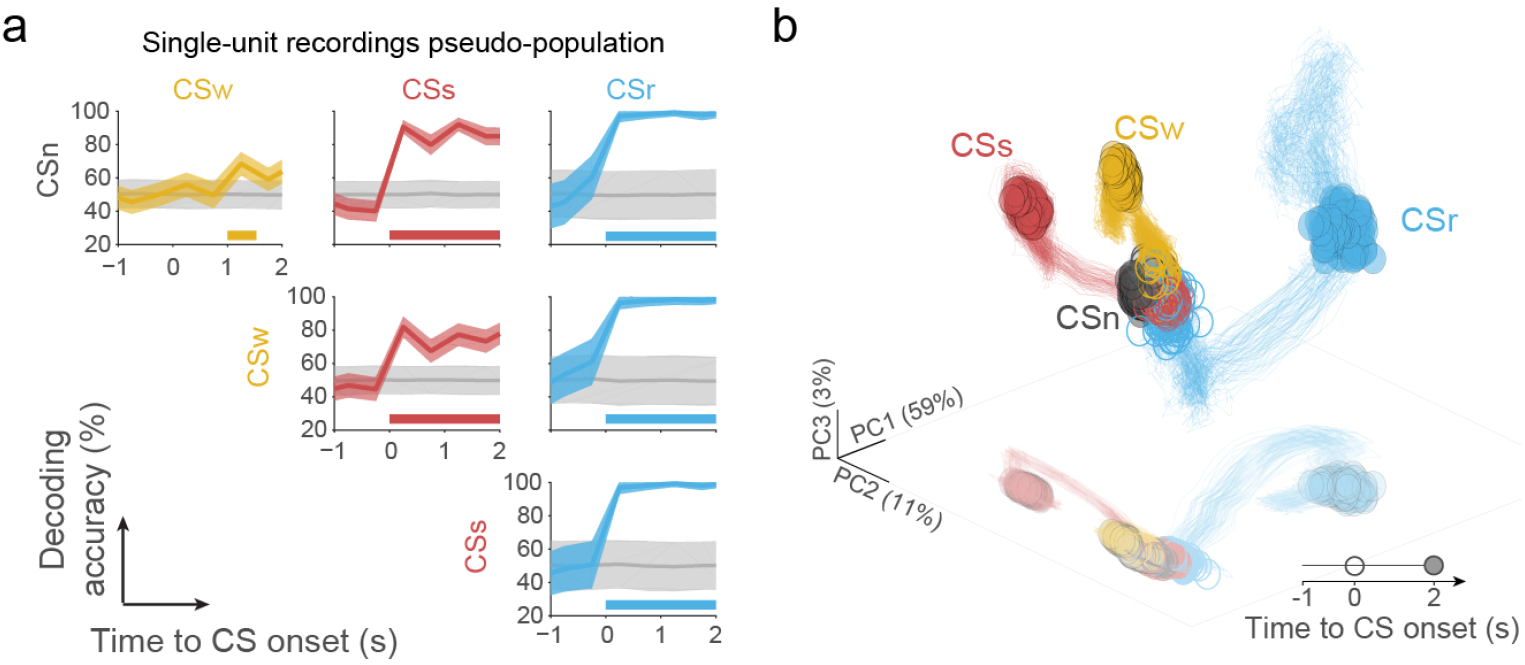
Electrophysiological single-unit recordings in the dmPFC during the approach/avoidance task. **a**. Decoding accuracies of different CS from dmPFC pseudo-population activity (n = 4 mice, 172 units). Accuracy data display mean +/- SD. Thick lines indicate significant decoding accuracies (P < 0.05, Permutation test). **b**. Neural trajectories for CS-onset triggered averaged responses for CSn, CSw, CSs, and CSr, projected onto the first 3 principal components (average responses over 80% of trials, 100 repetitions).

**Supplementary Figure 7.**
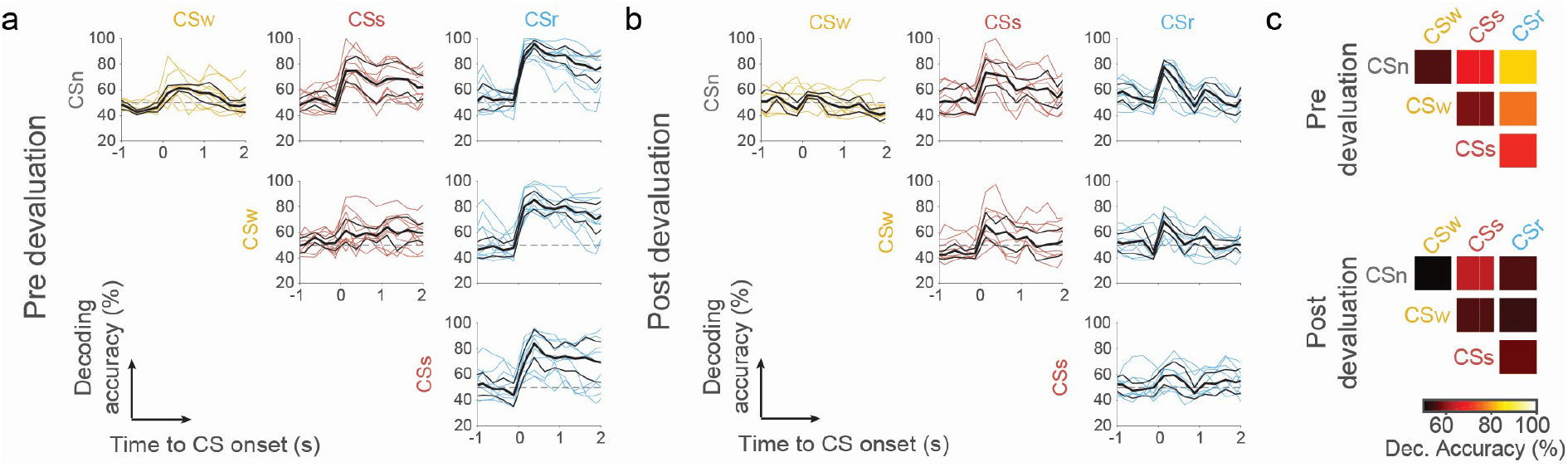
Impact of averaging activity across-cells (Multi single-cell) in decoding accuracies for single mice during the reward devaluation procedure (related to **Figure 3a-c**). **a**. Decoding accuracies Pre reward devaluation. **b**. Decoding accuracies Post reward devaluation. **c**. Median accuracy matrices for Pre (top) and Post (bottom) reward devaluation.

## Notes

### Competing Interest Statement

The authors have declared no competing interest.

